# Mosquito NF-κB-mediated innate immunity exerts arbovirus-specific antiviral effects at multiple stages of the viral life cycle

**DOI:** 10.1101/2025.11.06.687020

**Authors:** Philie Hollinghurst, Yin P. Cheung, Rayne Alexander, Tiffany A. Russell, Anthony C. Fredericks, Vikas Kumar, Louisa E. Wallace, Isabelle Dietrich, Tom A. Mendum, Andrew D. Davidson, Ana Fernandez-Sesma, Kevin Maringer

## Abstract

One third of all emerging infectious diseases are vector-borne, with the vector’s ecology and physiology playing key roles in determining whether viruses can access new vertebrate host species and spread globally. Innate immunity is a known barrier to virus replication in mosquito vectors that influences arboviral vector tropism. We here generated novel CRISPR-Cas9-mediated knockouts of the NF-κB family transcription factor Rel2 in *Aedes aegypti*-derived Aag2 cells and tested the impact on the replication of a diverse range of arboviruses in the *Flaviviridae* and *Togaviridae* families and the class *Bunyaviricetes*. We found that NF-κB-mediated innate immunity has broad antiviral activity against the *Ae. aegypti*-borne orthoflaviviruses dengue virus (DENV), yellow fever virus (YFV) and Zika virus (ZIKV) in mosquito cells. In contrast, little impact of NF-κB-loss-of-function was observed for the alphavirus chikungunya virus (CHIKV) or phlebovirus Rift Valley fever virus (RVFV), indicating specificity in the antiviral effects of NF-κB-mediated immunity. By comparing orthoflaviviruses with different transmission routes (mosquito-borne, tick-borne, no known vector), we demonstrated that NF-κB-mediated immunity exerts its antiviral effects both early and late in the viral replication cycle, and that NF-κB-mediated immunity is not the only molecular barrier influencing the ability of orthoflaviviruses to replicate in *Ae. aegypti* cells. Overall, our work demonstrates the importance of mosquito NF-κB-mediated innate immunity in suppressing arbovirus replication, and shows that the barriers for arboviruses to adapt to new vector species are multifactorial and virus-specific. Our findings increase our understanding of the molecular barriers influencing arboviral emergence, and could inform the development of refractory mosquitoes incapable of transmitting human pathogens.

**Author Summary:** Mosquito-borne viruses are causing an increasing global disease burden due to urbanisation, globalisation, changing land use and the climate change-driven invasion of mosquitoes into new locations. Few vaccines and no medicines for mosquito-borne viral infections are licensed for human use. Historically, the most effective way to reduce human disease has been to kill mosquitoes using insecticides, which also harms beneficial insects and is leading to resistance. Newer technologies with lower ecological impacts are being developed, including genetically modified mosquitoes with a reduced ability to transmit viruses. Like humans, mosquitoes have an immune system that protects against viral infections, and strengthening this immune system through genetic modification has shown promise for reducing virus transmission. We here expand our understanding of the ways in which mosquito immunity could be harnessed by demonstrating that a gene called “Rel2” helps the mosquito fight mosquito-borne virus infection. Unlike previously studied mosquito immune responses, Rel2 has a very broad (but not universal) protective capacity against viruses. Our work also indicates that Rel2 works in concert with other arms of the mosquito immune system, highlighting a need for further research to fully realise the potential of genetically modifying mosquito immune responses for the prevention of human disease.

## Introduction

One third of all emerging infectious diseases are vector-borne [1], with the vector’s ecology and physiology playing key roles in determining whether viruses can access new vertebrate host species or geographic regions [2,3]. The most impactful arthropod-borne viruses (arboviruses) affecting humans are in the *Flaviviridae* and *Togaviridae* families and the class *Bunyaviricetes*. However, the molecular barriers that influence the vector tropism of these arboviruses (the ability of a virus to replicate in one vector over another and hence access one or other vertebrate host species) remain incompletely understood.

*Aedes aegypti* is the most significant vector of arthropod-borne viruses (arboviruses) of humans, and understanding the molecular determinants influencing its ability to transmit arboviruses (“vector competence”) is crucial to reducing the global burden of arboviral disease. In the absence of antiviral therapies and with few effective vaccines available, vector control strategies have remained the most viable means of protecting people from arboviral infection. Traditional vector control strategies have relied on the use of insecticides, which, though effective in achieving population suppression, elicit off-target ecological consequences and select for resistance. Population modification to reduce the capacity of the vector to transmit arboviruses, either through genetic modification or the introgression of the arbovirus-restricting intracellular bacterium *Wolbachia pipientis*, presents an attractive alternative with proven success in the lab and field [4–9].

Mosquito immunity poses a significant endogenous barrier to the replication and transmission of arboviruses by mosquitoes [10]. The most well-studied immune responses of mosquitoes are RNA interference (RNAi) pathways, sequence-specific antiviral responses mediated by virus-restricting small interfering RNAs (siRNAs) and piwi-interacting RNAs (piRNAs) [11,12]. Refractory *Ae. aegypti* with reduced vector competence for the orthoflaviviruses dengue virus (DENV, *Orthoflavivirus denguei*) and Zika virus (ZIKV, *Orthoflavivirus zikaense*) have been generated by exploiting RNAi-mediated silencing of viral RNA using transgenically expressed small RNA constructs [9,13]. Inducible non-sequence-specific immune responses, in which pathogen detection activates a signalling cascade leading to the expression of broadly active antimicrobial genes, remain less well characterised in the context of viral infection but might offer broader protection against a wider range of unrelated arboviruses.

The nuclear factor kappa-light-chain-enhancer of activated B cells (NF-κB) family of transcription factors is highly conserved across vertebrates and invertebrates and regulates the expression of antimicrobial genes in mosquitoes [14]. *Ae. aegypti* encodes several NF-κB family transcription factors. Rel1A acts in concert with Rel1B and is canonically thought of as being activated via Toll-like receptors (TLRs) within the Toll pathway (Fig 1A) [14]. Rel2 meanwhile is canonically thought of as being activated through peptidoglycan recognition proteins (PGRPs) via the immunodeficiency (Imd) protein as part of the Imd pathway (Fig 1A) [14]. We [15] and others [16,17] showed that NF-κB-regulated genes can be induced by the viral double-stranded RNA (dsRNA) mimic poly(I:C) as well as viral infection in mosquitoes *in vivo* and in cell culture. Furthermore, stimulating NF-κB signalling using poly(I:C) or bacteria reduces the replication of DENV [16] and the alphavirus Semliki Forest virus (SFV, *Alphavirus semliki*, family *Togaviridae*) [18] respectively. However, exogenous immune stimulation has the potential to activate multiple pathways, and the gold standard for demonstrating the antiviral potential of an innate immune pathway is therefore genetic manipulation of the pathway itself. Transiently silencing mosquito NF-κB-inducing pattern recognition receptors (PRRs) or NF-κB-regulated antimicrobial peptides (AMPs) results in enhanced arboviral replication *in vivo* and in cell culture [17,19–22]. However, PRRs and AMPs exhibit crosstalk between multiple NF-κB-regulated innate immune pathways [23–28], and conclusions on the specificity of the antiviral effects of a particular NF-κB-regulated pathway are difficult to draw from these data.

**Figure 1.**
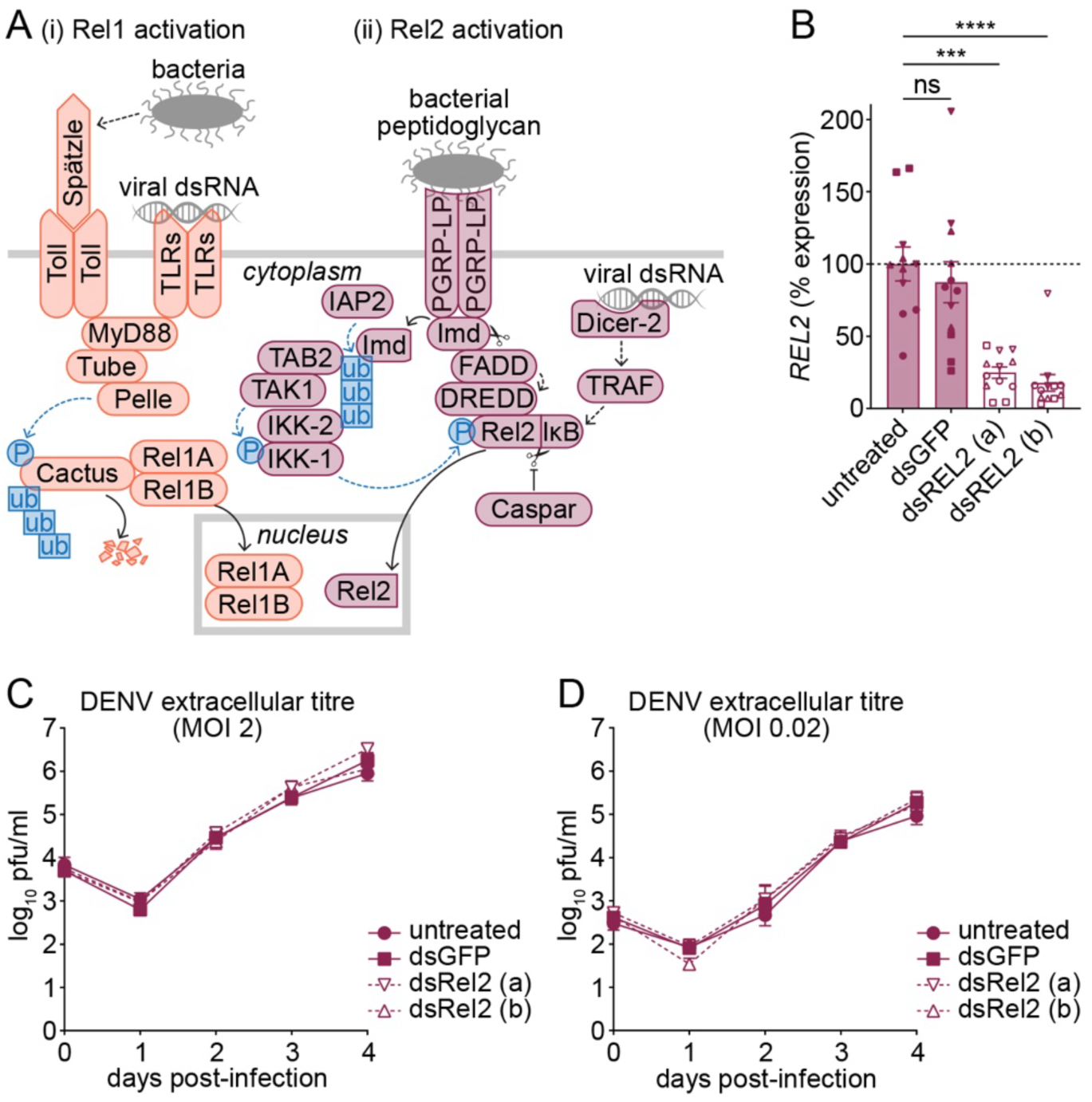
Transient Rel2 knockdown has no effect on DENV-2 replication in Aag2-AF5 cells. (A) Summary of mosquito NF-κB pathways. (i) Rel1 is activated when Toll and Toll-like receptors (TLRs) sense viral or bacterial infection, in the case of bacteria via the proteolytic activation of the cytokine Spätzle. The subsequent recruitment of myeloid differentiation primary response 88 (MyD88), Tube and Pelle leads to the phosphorylation, ubiquitination and proteasomal degradation of the negative regulator Cactus, allowing a complex of the NF-κB family transcriptional regulators Rel1A and Rel1B to translocate into the nucleus and activate antimicrobial gene expression. (ii) Rel2 is activated following detection of viral dsRNA by Dicer-2, through an incompletely characterised signalling cascade involving tumour necrosis factor (TNF)-associated factor (TRAF). In addition, bacterial peptidoglycan is detected by peptidoglycan recognition proteins (e.g. PGRP-LP), leading to recruitment and proteolytic cleavage of the immunodeficiency protein (Imd) and subsequent K63 ubiquitination-mediated activation of the inhibitor of κB (IκB) kinase (IKK) complex by the inhibitor of apoptosis family protein IAP2, transforming growth factor-β-activated kinase 1 (TAK1) and TAK1-binding protein 2 (TAB2). The Imd-dependent recruitment of Fas-associated protein with death domain (FADD) and death-related ced-3/Nedd2-like protein (DREDD) also helps facilitate the phosphorylation and proteolytic cleavage of the NF-κB family transcription factor Rel2, releasing its C-terminal inhibitory IκB domain and allowing its translocation to the nucleus to activate antimicrobial gene expression. Caspar is a negative regulator of Rel2 cleavage. (B) RT-qPCR measuring Rel2 transcript levels in Aag2-AF5 cells following transient transfection with two independent non-overlapping dsRNAs targeting Rel2 (dsREL2), or an irrelevant control dsRNA specific for green fluorescent protein (dsGFP). (C and D) DENV-2 replication in Aag2-AF5 cells following Rel2 knockdown and infection at an MOI of 2 (C) or 0.02 (D), as measured by titration of extracellular infectious particles in plaque-forming units (pfu)/ml. Data represent mean ± standard error of the mean (SEM); minimum *N* = 3, *n* = 3; in panel B, data point shapes indicate biological replicates (i.e. same shape indicates technical replicates within one experiment). Unless indicated, differences are not significant; *** *P* < 0.001; **** *P* < 0.0001; ns, not significant.

In this study we aimed to establish whether Rel2-mediated innate immunity is antiviral in *Ae. aegypti* and to define the extent to which Rel2-mediated immunity presents a barrier to vector switching in arboviruses. To this end, we generated novel CRISPR-Cas9-mediated knockouts of *REL2* in *Ae. aegypti*-derived Aag2 cells and tested the impact on the replication of a diverse range of arboviruses in the *Flaviviridae* and *Togaviridae* families and the class *Bunyaviricetes*. Rel2 was found to restrict a broad range of orthoflaviviruses, but not the alphavirus chikungunya virus (CHIKV, *Alphavirus chikungunya*) or phlebovirus Rift Valley fever virus (RVFV, *Phlebovirus riftense*). Furthermore, we demonstrated that Rel2 exerts antiviral effects both early and late in the viral replication cycle. Notably, Rel2 knockout did not rescue replication of mosquito-restricted orthoflaviviruses, indicating that additional barriers to vector switching exist in *Ae. aegypti*. We therefore present the first direct evidence for broad antiviral properties of a specific NF-κB-regulated pathway in mosquitoes, and identify complexity in the mechanisms by which NF-κB-mediated immunity restricts the replication cycle of arboviruses. These findings increase our understanding of the molecular barriers influencing arboviral emergence, and could inform the development of refractory mosquitoes incapable of transmitting human pathogens.

## Results

### Transiently silencing Rel2 has no effect on DENV-2 replication in *Ae. aegypti* cells

We first attempted to investigate the antiviral activity of Rel2-mediated innate immunity by transiently knocking down Rel2 expression in *Ae. aegypti*-derived Aag2-AF5 cells using two independent non-overlapping dsRNAs, with an irrelevant dsRNA targeting green fluorescent protein (GFP) serving as a negative control. Having confirmed Rel2 knockdown (Fig 1B), we tested its impact on the replication of the orthoflavivirus DENV-2. However, we saw no significant difference in the accumulation of extracellular infectious DENV-2 particles in the cell culture supernatant of Rel2-depleted Aag2-AF5 cells compared to controls when cells were infected at a multiplicity of infection (MOI) of 2 (Fig 1C). Since antiviral effects can be more pronounced at low MOIs due to slower rates of virus replication that are more readily restricted by the immune system [29], we also tested the impact of Rel2 knockdown at an MOI of 0.02, and again observed no statistically significant difference in extracellular infectious particle accumulation compared to control cells (Fig 1D). Taken together, these data show that the *Ae. aegypti* Rel2 signalling pathway does not exert overt antiviral effects against DENV-2 when Rel2 expression is transiently silenced.

### Generation of an *Ae. aegypti*-derived Rel2 knockout cell line using CRISPR-Cas9-mediated gene editing

We hypothesised that Rel2’s antiviral effects would be more pronounced following gene knockout using CRISPR-Cas9-mediated gene editing, and therefore generated a clonal Rel2 knockout cell line using our previously established protocols [30,31]. The Rel2 knockout cell line (which we termed Aag2-AF256) is derived from Aag2-AF5 cells, a clonal cell line that we previously derived from the heterogenous Aag2 cell line to minimise the impact of clonal bottlenecks during knockout cell line generation [32].

### Rel2 elicits antiviral effects against a broad range of orthoflaviviruses

Next, we repeated our DENV-2 infections at high and low MOIs in Aag2-AF256 cells and compared viral replication with wild-type Aag2-AF5 cells (Fig 3A, 3B). Similarly to our data from transient silencing of Rel2 (Fig 1C), at an MOI of 2, the accumulation of extracellular infectious DENV-2 particles was comparable in Aag2-AF5 and Aag2-AF256 cells (Fig 3Ai). Furthermore, no difference in the accumulation of intracellular DENV-2 RNA was observed in Rel2 knockout cells compared to wild-type cells at this MOI (Fig 3Aii). However, when we repeated the infections at an MOI of 0.02, we did observe a modest (less than one log) but statistically significant increase in the maximum extracellular titre of DENV-2 late in infection (day 6) in Aag2-AF256 cells compared to Aag2-AF5 cells (Fig 3Bi), demonstrating that loss of functional Rel2 results in reduced antiviral activity in *Ae. aegypti*-derived cell cultures. The accumulation of intracellular DENV-2 RNA was not elevated in Rel2 knockout versus wild-type cells up to three days post-infection (Fig 3Bii).

**Figure 3.**
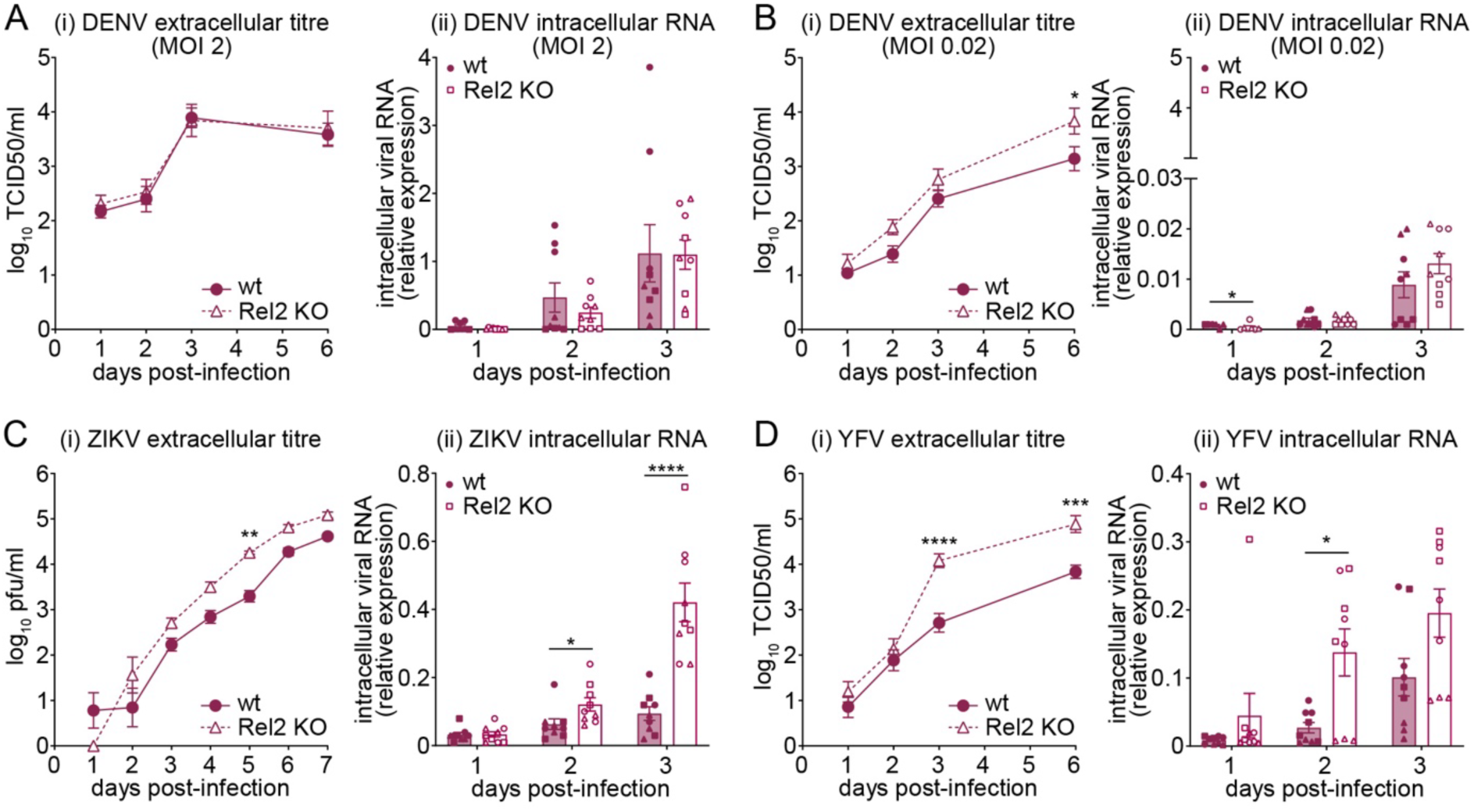
Rel2 is antiviral against *Aedes*-borne orthoflaviviruses. (A) Replication of DENV-2 in Aag2-AF5 (wt) or Aag2-AF256 (Rel2 KO) cells following infection at MOI 2. (B-D) Replication of DENV-2 (B), ZIKV (C) or YFV (D) in Aag2-AF5 (wt) or Aag2-AF256 (Rel2 KO) cells following infection at MOI 0.02. (i) Extracellular infectious particles quantified by TCID50 (TCID50/ml) or plaque assay (pfu/ml) as indicated. (ii) Intracellular viral RNA quantified by RT-qPCR and expressed relative to *RPS7* housekeeping gene (2^-ΔCt^); for Bii relative expression represents 100 × 2^-ΔCt^. Data represent mean ± SEM; *N* = 3, *n* = 3; in all (ii) panels, data point shapes indicate biological replicates (i.e. same shape indicates technical replicates within one experiment). Unless indicated, differences are not significant; * *P* < 0.05; ** *P* < 0.01; *** *P* < 0.001; **** *P* < 0.0001; ns, not significant.

We next explored how broad the antiviral effects of Rel2 are against other orthoflaviviruses transmitted by *Ae. aegypti*, repeating the experiment with ZIKV and yellow fever virus (YFV, *Orthoflavivirus flavi*). Since the antiviral effects of Rel2 were only observed at low MOI, these and all subsequent viral infections were performed at an MOI of 0.02. The accumulation of extracellular infectious ZIKV particles was elevated by up to one-log across all time points post-infection in Aag2-AF256 cells compared to Aag2-AF5 cells, with extracellular infectivity reaching comparable levels one day faster than in wild-type cells (Fig 3Ci). Intracellular ZIKV RNA levels were also significantly elevated in Rel2 knockout cells compared to wild-type cells; up to 4.5-fold higher by three days post-infection (Fig 3Cii). Similarly, YFV replication was significantly higher in Rel2 knockout versus wild-type cells, with up to two-logs higher extracellular infectivity and up to five-fold higher intracellular RNA levels observed in Aag2-AF256 cells compared to Aag2-AF5 cells (Fig 3D). Taken together, our data identify broad antiviral effects of Rel2-mediated innate immune signalling against a range of *Ae. aegypti*-borne orthoflaviviruses, and these antiviral effects are already observed early in infection prior to the accumulation of viral RNA.

### Rel2 is not antiviral against a representative alphavirus and bunyavirus

We next explored how broadly effective the antiviral action of Rel2-mediated immunity is on unrelated arboviruses transmitted by *Aedes spp.* mosquitoes. Interestingly, we found no evidence of Rel2-mediated suppression of the alphavirus CHIKV, with no significant differences in viral replication observed in Aag2-AF256 versus Aag2-AF5 cells at the level of intracellular viral RNA or extracellular infectious particle accumulation (Fig 4A). Similarly, levels of extracellular infectious particles and intracellular viral RNA were unaffected by Rel2 knockout when cells were infected with the phlebovirus RVFV (Fig 4B). Collectively, our data therefore indicate a degree of specificity in the antiviral effects of Rel2, which appears to primarily act on flaviviruses, but not alphaviruses or bunyaviruses, at least under the conditions tested in our cell culture experiments.

**Figure 4.**
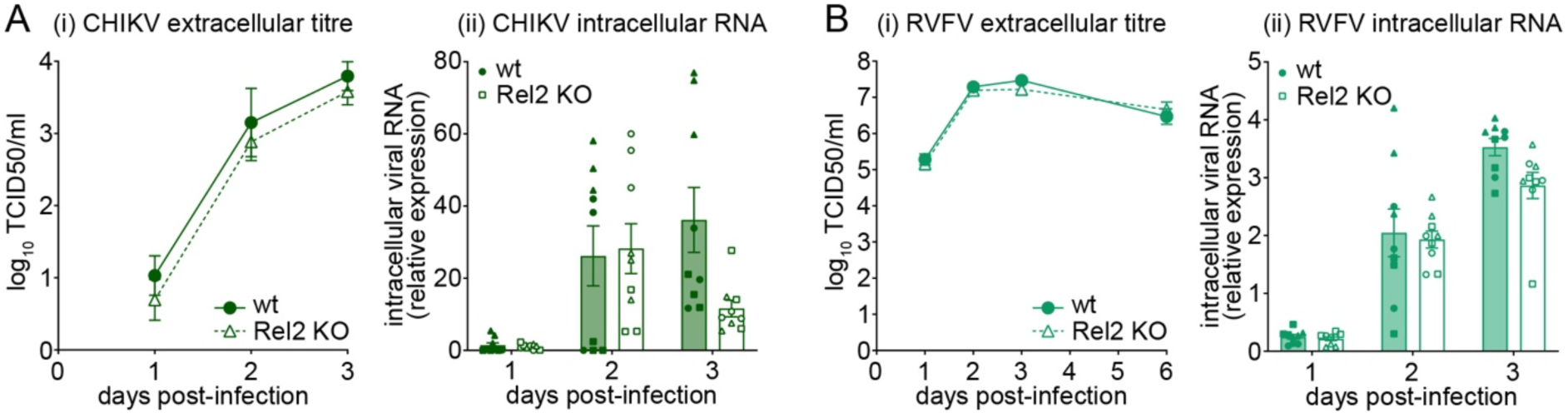
Rel2 is not antiviral against a representative alphavirus and bunyavirus. Replication of CHIKV (A) or RVFV (B) in Aag2-AF5 (wt) or Aag2-AF256 (Rel2 KO) cells following infection at MOI 0.02. (i) Extracellular infectious particles quantified by TCID50 (TCID50/ml). (ii) Intracellular viral RNA quantified by RT-qPCR and expressed relative to *RPS7* housekeeping gene (2^-ΔCt^). Data represent mean ± SEM; *N* = 3, *n* = 3; in all (ii) panels, data point shapes indicate biological replicates (i.e. same shape indicates technical replicates within one experiment). Differences between wild-type and Rel2 knockout cells are not statistically significant at any time point.

### Rel2-mediated innate immunity has a virus-specific antiviral effect among mosquito-borne orthoflaviviruses

To further explore the extent to which Rel2-mediated innate immunity exerts virus-specific antiviral effects, we tested additional mosquito-borne orthoflaviviruses. Both West Nile virus (WNV, *Orthoflavivirus nilense*) and Japanese encephalitis virus (JEV, *Orthoflavivirus japonicum*) are transmitted primarily by *Culex spp.* mosquitoes, but can establish experimental infection in *Ae. aegypti* and *Ae. aegypti*-derived cell cultures in the lab. Similarly to the other orthoflaviviruses tested, the accumulation of extracellular infectious WNV particles was elevated, by up to two logs, in Rel2 knockout versus wild-type cells (Fig 5Ai). However, in contrast to other orthoflaviviruses, no differences in the accumulation of intracellular WNV RNA were observed following Rel2 knockout (Fig 5Aii), indicating that as well as exerting antiviral effects early in the orthoflavivirus replication cycle that influence the accumulation of viral RNA (Fig 3), Rel2-mediated immune responses also target later stages of the viral life cycle.

**Figure 5.**
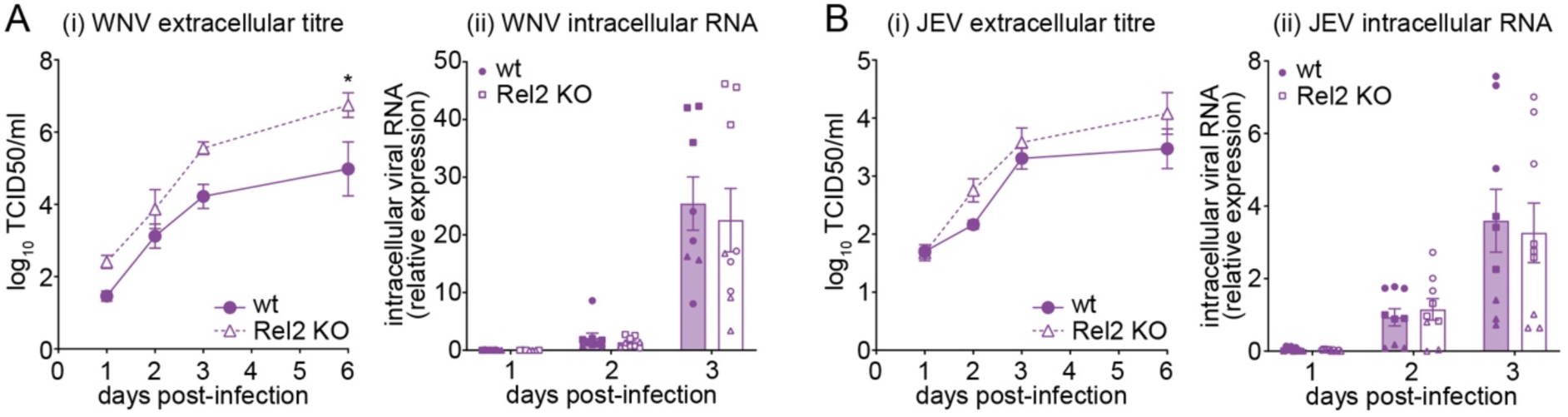
Rel2 exerts antiviral effects on *Culex*-borne orthoflaviviruses late in the replication cycle. Replication of WNV (A) or JEV (B) in Aag2-AF5 (wt) or Aag2-AF256 (Rel2 KO) cells following infection at MOI 0.02. (i) Extracellular infectious particles quantified by TCID50 (TCID50/ml). (ii) Intracellular viral RNA quantified by RT-qPCR and expressed relative to *RPS7* housekeeping gene (2^-ΔCt^). Data represent mean ± SEM; *N* = 3, *n* = 3; in all (ii) panels, data point shapes indicate biological replicates (i.e. same shape indicates technical replicates within one experiment). Unless indicated, differences are not significant; * *P* < 0.05.

Although we observed a trend towards elevated levels of extracellular infectious particle accumulation for JEV in Aag2-AF256 compared to Aag2-AF5 cells, none of the observed differences were statistically significant (Fig 5Bi). Furthermore, no difference in the accumulation of intracellular viral RNA was observed for JEV in Rel2 knockout compared to wild-type cells (Fig 5Bii). We therefore conclude that Rel2-mediated innate immunity exerts antiviral effects that are not only virus-specific at the viral family level, but also within the *Orthoflavivirus* genus.

### Rel2 is not the only molecular barrier to orthoflavivirus replication in *Ae. aegypti* cells

In light of our finding that Rel2-mediated innate immunity is antiviral against a broad range of orthoflaviviruses, we hypothesised that Rel2 may constitute an important component of the molecular barrier to orthoflavivirus vector-switching. We therefore infected Aag2-AF5 and Aag2-AF256 cells with tick-borne orthoflaviviruses that cannot replicate in *Ae. aegypti* or *Ae. aegypti*-derived cells. When cells were infected with Langat virus (LGTV, *Orthoflavivirus langatense*), infectious particles introduced with the inoculum persisted in both Rel2-knockout as well as wild-type cells but no increase in extracellular infectivity indicative of viral replication was observed over time (Fig 6Ai). Similarly, no accumulation of intracellular LGTV RNA was detected in either Aag2-AF5 or Aag2-AF256 cells (Fig 6Aii). Importantly, we confirmed the infectivity of our LGTV stock by infecting mammalian Vero cells, in which replication was observed as measured by an increase in the amount of extracellular infectious particles (Fig 6Aiii). We corroborated this finding with another tick-borne orthoflavivirus, Powassan virus (POWV, *Orthoflavivirus powassanense*), for which we also failed to detect any extracellular infectious particle or intracellular viral RNA accumulation in either Rel2 knockout or wild-type cells, despite confirming the infectivity of our viral stock in Vero cells (Fig 6B).

**Figure 6.**
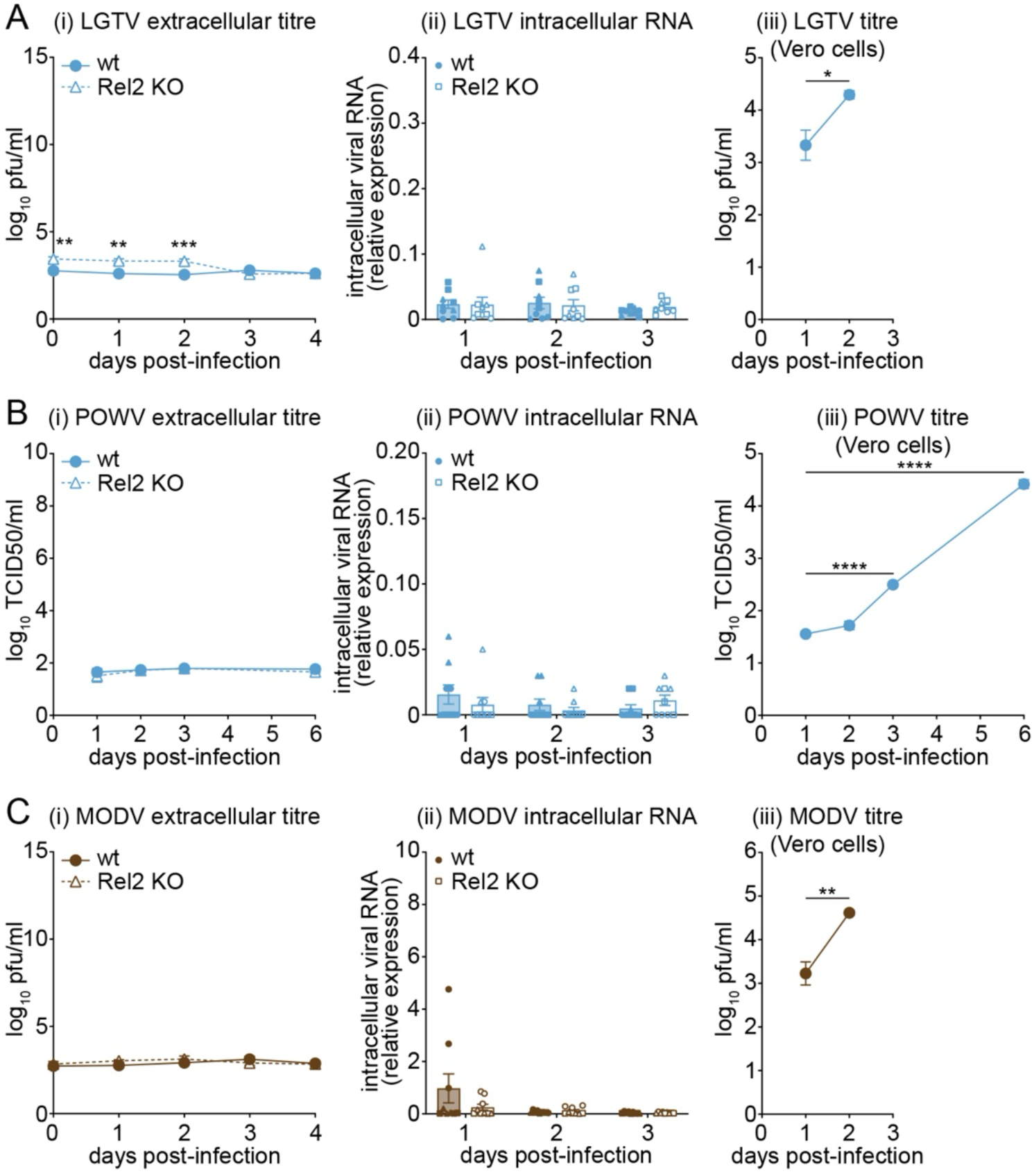
Rel2 is not the only molecular barrier restricting orthoflavivirus replication in *Ae. aegypti* cells. Replication of LGTV (A), POWV (B) or MODV (C) in Aag2-AF5 (wt) and Aag2-AF256 (Rel2 KO) cells (i, ii) or Vero cells (iii) following infection at MOI 0.02. (i, iii) Extracellular infectious particles quantified by TCID50 (TCID50/ml) or plaque assay (pfu/ml) as indicated. (ii) Intracellular viral RNA quantified by RT-qPCR and expressed relative to *RPS7* housekeeping gene (2^-ΔCt^). Data represent mean ± SEM; *N* = 3, *n* = 3; in all (ii) panels, data point shapes indicate biological replicates (i.e. same shape indicates technical replicates within one experiment). Unless indicated, differences are not significant; * *P* < 0.05; ** *P* < 0.01; *** *P* < 0.001; **** *P* < 0.0001.

Finally, we also tested the no known vector orthoflavivirus Modoc virus (MODV, *Orthoflavivirus modocense*), which also does not replicate in mosquito cells. Similarly to the tick-borne viruses that we tested, MODV did not replicate in either Aag2-AF5 or Aag2-AF256 cells, with no accumulation of extracellular infectious particles or intracellular RNA observed, but was able to replicate in Vero cells (Fig 6C). The lack of replication of LGTV, POWV and MODV suggests that while Rel2-mediated immunity is antiviral against orthoflaviviruses, additional molecular barriers to replication in *Ae. aegypti*-derived cells exist for viruses not adapted to mosquito-borne transmission.

## Discussion

In this study, we generated a novel Rel2 knockout cell line in *Ae. aegypti*-derived Aag2 cells, and present evidence for the antiviral activity of NF-κB against a broad range of orthoflaviviruses in *Ae. aegypti*. We were only able to demonstrate Rel2’s antiviral effects against DENV-2 following complete genetic knockout of Rel2. Similarly, ZIKV replication was also not enhanced when Rel2 and other signalling pathway components leading to Rel2 activation were transiently silenced in *Ae. aegypti in vivo* [21], indicating that some antiviral functions may only be identifiable through complete genetic knockout. We found the antiviral effects of Rel2 to be selective, with no observed impact on the replication of a representative alphavirus (CHIKV) or bunyavirus (RVFV). Additionally, in our hands, Rel2 knockout did not rescue replication of mosquito-restricted orthoflaviviruses in Aag2 cells, suggesting that additional molecular barriers influencing the species-specificity of orthoflaviviruses exists.

### On the antiviral nature of *Ae. aegypti* NF-κB-mediated immunity against orthoflaviviruses

In *Ae. aegypti*, the antiviral nature of the Rel2 signalling pathway against orthoflaviviruses had previously only been demonstrated indirectly through the transient knockdown of upstream PRRs or downstream NF-κB-regulated AMPs [19,22,34], with the interpretation of these data confounded by crosstalk between NF-κB-regulated innate immune pathways [23–28]. Furthermore, there are complex interactions between NF-κB signalling pathways and the insect microbiome that jointly influence arbovirus replication [35–38]. NF-κB signalling also modulates peritrophic matrix formation *in vivo*, which modifies susceptibility to arboviral infection [39–41]. It is therefore significant that we here demonstrate antiviral effects of Rel2 against orthoflaviviruses in cell culture in the absence of these complex interactions, proving that Rel2 can elicit direct cell-intrinsic antiviral effects.

In *Drosophila melanogaster*, cyclic GMP-AMP synthase (cGAS)-like receptor (cGLR)-stimulator of interferon genes (STING) signalling, which signals into the Rel2 homologue Relish, is antiviral [42–45]. However, in this model insect, there is no genetic evidence for antiviral effects mediated by Relish beyond signalling pathway components also implicated in cGLR-STING signalling [14]. cGLR-STING signalling is not conserved in mosquitoes [46], and our data therefore demonstrate either that the canonical Imd pathway signalling into Rel2 is antiviral, or that additional virus-specific pathways leading to Rel2 activation exist in mosquitoes, and perhaps other insects. While some potential PRRs that sense viral infection to activate NF-κB family transcription factors have been identified in *Aedes spp.* and *Culex spp.* mosquitoes, namely TLR6, scavenger receptors and Dicer-2 [16,17,19,47], their role in antiviral NF-κB-mediated immunity and the precise signalling cascades they activate remain to be fully elucidated. Furthermore, antiviral effects have been shown for some NF-κB-regulated defensins and cecropins that were originally identified as antibacterial [19,34], though additional virus-specific NF-κB-regulated genes likely remain to be discovered. Our findings provide renewed vigour and rationale for a concerted effort to characterise in detail the virus-specific signalling cascades leading to NF-κB activation in insects, as well as the NF-κB-regulated downstream effectors that are specifically antiviral.

### Complexity in the NF-κB-mediated antiviral response

Two of our findings hint at hitherto unappreciated complexity in the NF-κB-mediated antiviral response. Firstly, we found that Rel2 knockout resulted in higher DENV-2 replication at low, but not high, MOI. This may be related to DENV’s ability to dampen NF-κB-regulated gene expression in *Ae. aegypti* cells and *in vivo* [48–50], suggestive of the existence of virally encoded immune antagonists allowing the virus to outcompete the Rel2-dependent antiviral response at higher MOIs when viral proteins accumulate more rapidly and at higher levels. Alternatively, it may be that Rel2’s antiviral effects are predominantly exerted in trans through cytokine-like signals that establish an antiviral state in bystander cells, in which case any associated antiviral effects would be more pronounced at lower MOIs when cell-to-cell viral spread becomes more important. In line with this hypothesis, “interferon-like” cytokines in the Vago family, which in some insects have been shown to stimulate the antiviral Janus kinase-signal transducer and activator of transcription (Jak-STAT) pathway [51,52], are regulated by Rel2 in *Culex quinquefasciatus*-derived Hsu cells [47]. However, recent *in vivo* data from *Ae. aegypti* raise questions about the interpretation of earlier findings [53], and we failed to detect Vago-like gene expression in Aag2 cells [15], indicating that more research is needed in this area. Finally, infection with the same virus at high versus low MOIs has been shown to induce different NF-κB-regulated antiviral genes, and through different mechanisms, in mammals [54], and it is feasible that MOI similarly modulates NF-κB-regulated antiviral responses in mosquitoes.

Our second interesting finding is that Rel2 exerts its effects at both early and late stages of the viral replication cycle. For most of the orthoflaviviruses tested, enhanced replication in Rel2 knockout cells was already observed at the RNA level, indicating that Rel2 exerts antiviral effects during viral entry, uncoating, translation and/or genome replication. In contrast, for WNV, levels of extracellular infectious particles were elevated without a concomitant increase in intracellular viral RNA levels, indicating that Rel2 can also exert antiviral effects later in infection, affecting virion assembly, release and/or maturation. Little is known about the mechanisms by which NF-κB-regulated genes exert their antimicrobial effects in insects, and these molecular details will be important to clarify in future studies.

### Specificity of Rel2’s antiviral effects

In our hands, Rel2 was broadly antiviral against orthoflaviviruses, but had no effect on the alphavirus CHIKV or the bunyavirus RVFV. Taken together with other published studies however, a clear picture does not emerge on the antiviral role of mosquito Rel2 on alphaviruses and bunyaviruses. While some studies did observe enhanced replication of CHIKV in Aag2 cells [55] and O’nyong-nyong virus (ONNV, *Alphavirus onyong*) in *Anopheles gambiae in vivo* [56] following Rel2 knockdown, other studies did not replicate these findings [20,57]. Meanwhile, our RVFV data agrees with studies indicating that RVFV infection does not by itself induce NF-κB-regulated genes in Aag2 cells [58]. In contrast, RVFV did induce NF-κB-regulated genes in *C. pipiens in vivo* while La Crosse virus (LACV, *Orthobunyavirus lacrosseense*) replication was found to be enhanced in *C. quinquefasciatus*-derived Hsu cells following transient silencing of Rel2 [17,59]. There is therefore a need for larger systematic studies exploring the antiviral roles of Rel2 against alphaviruses and bunyaviruses in different vector species.

Furthermore, despite exerting broadly antiviral effects against most of the orthoflaviviruses tested, JEV was not restricted by Rel2, demonstrating a level of specificity in Rel2’s antiviral activity within the orthoflavivirus genus. Intriguingly, although typically antiviral [19,34], some defensins enhance JEV binding to *Culex pipiens pallens* and *Aedes albopictus* cells [60]. The literature also presents conflicting evidence on whether signalling into Rel2 is antiviral or proviral in the context of ZIKV infection in *Ae. aegypti in vivo* [20,21], while Rel2 knockdown in *C. quinquefasciatus*-derived Hsu cells corroborates Rel2’s antiviral effect against WNV [47]. Overall, these findings indicate that NF-κB pathways may exert a complex concert of both antiviral and proviral roles, with responses being potentially virus- and mosquito species-specific.

### NF-κB-mediated immunity as a determinant of arboviral vector tropism

In our hands, tick-borne and no-known-vector orthoflaviviruses did not productively replicate in *Ae. aegypti* cells in the presence or absence of Rel2, indicating that additional molecular barriers influencing arboviral vector-specificity exist. Host range restriction of orthoflaviviruses has previously been shown to occur post-entry [61–63], consistent with innate immune responses and other post-entry barriers determining the vector tropism of orthoflaviviruses. There is evidence suggesting that activation of Rel1A via MyD88 is also antiviral against DENV [64,65]. Furthermore, LGTV has been shown to replicate in *Ae. albopictus*-derived C6/36 cells, which are RNAi-deficient [66], potentially suggesting that RNAi influences arboviral vector tropism. On the other hand, POWV failed to replicate in C6/36 cells [67], potentially indicative of virus-specific restriction of arbovirus replication by vector immunity. Either way, the role of RNAi and other immune responses in restricting arboviruses in their non-cognate vectors remains to be rigorously tested and compared to determine the relative importance of individual pathways. Interestingly, the relative enhancement of viral replication in our Rel2 knockout and previously published RNAi-deficient cell lines is comparable [30,31,68], indicating that at a cell intrinsic level the RNAi pathway is not overtly more restrictive to viral replication than NF-κB-mediated immunity. It will now be important to directly compare the relative antiviral impacts of RNAi and NF-κB pathways, and explore whether they act in concert or separately to restrict arbovirus replication and transmission in cell culture and *in vivo*.

In summary, our data provide important new evidence for the antiviral role of NF-κB-mediated immunity in mosquitoes, hinting at complex NF-κB-regulated antiviral effectors that block arboviral replication at multiple stages of the viral life cycle, and demonstrating a degree of specificity towards restriction of orthoflaviviruses. It will be interesting in the future to explore whether and how NF-κB-mediated immunity could be exploited to develop refractory mosquitoes less able to transmit arboviruses, given that the transgenic expression of AMPs has been shown to restrict *Plasmodium spp.* transmission in *An. gambiae* [69]. It will also be important to test the impact of Rel2 knockout *in vivo* given that the potent and reproducible antiviral effects of RNAi-mediated immunity in cell culture [30,31,68] are less pronounced *in vivo* [70–72]. Overall, our data expand our understanding of the molecular factors influencing vector competence and arboviral vector tropism at a time when the global impact of arboviruses is increasing due to globalisation, urbanisation, and climate and land-use change.

## Materials and Methods

### Cells

Aag2-AF5 cells are a clonal derivative of Aag2 cells that we previously generated, available through the European Collection of Authenticated Cell Cultures (ECACC, culturecollections.org.uk, catalogue number 19022601) [32].

Aag2 cells and derived clones were cultured in Leibovitz’s L-15 medium (Sigma-Aldrich, St. Louis, MO USA) supplemented with 10% (v/v) tryptose phosphate broth (Sigma-Aldrich), 10% (v/v) foetal bovine serum (FBS), 2 mM L-glutamine, 0.1 mM non-essential amino acids, 100 U/ml penicillin and 100 μg/ml streptomycin (all ThermoFisher Scientific). Medium for initial expansion of single cell clones contained 20% (v/v) FBS. Cells were grown at 28°C in a humidified atmosphere without CO_2_.

Mammalian baby hamster kidney (BHK)-21, Vero and Vero E6 cells were sourced from ECACC with respective catalogue numbers 85011433, 84113001 and 85020206. Cells were cultured in high glucose Dulbecco’s modified Eagle’s medium (DMEM) containing pyruvate and supplemented with 10% (v/v) FBS, 100 U/ml penicillin and 100 μg/ml streptomycin (all ThermoFisher Scientific). Cells were grown at 37°C in a humidified atmosphere with 5% CO_2_. C6/36 cells were a kind gift from Jorge Muñoz-Jordan (Centers for Disease Control and Prevention, San Juan, Puerto Rico), and were cultured in Roswell Park Memorial Institute (RPMI) medium supplemented with 0.15% (w/v) sodium bicarbonate (Sigma-Aldrich), 0.1 mM non-essential amino acids, 2 mM L-glutamine, 1 mM sodium pyruvate and 10% (v/v) FBS (all ThermoFisher Scientific) at 33°C in a humidified atmosphere with 5% CO_2_. All cell lines were regularly confirmed to be free of mycoplasma contamination.

### Viruses

DENV-2 strain 16681 and LGTV strain TP21 were kind gifts from Ana Fernandez-Sesma and Jean Lim respectively (Icahn School of Medicine at Mount Sinai, New York, NY USA). ZIKV strain Paraiba_01 was a kind gift from Stephen Whitehead (National Institute of Allergy and Infectious Diseases, Bethesda, MD USA). MODV strain M544 was a kind gift from Zahra Zakeri (Queens College, New York, USA). POWV strain POW/2 and WNV strain NY99 were kind gifts from Karen Mansfield (Animal and Plant Health Agency, Addlestone, UK). Recombinant RVFV strain ZH548 were kind gifts from Isabelle Dietrich (The Pirbright Institute). CHIKV strain LR2006OPY1 was a kind gift from Christine Reitmayer (Keele University, Keele, UK). JEV strain CNS769/Laos/2009 (genotype I) and YFV strain Asibi were obtained through the European Virus Archive global (EVAg, european-virus-archive.com).

For CHIKV, JEV, LGTV, MODV, POWV, RVFV, WNV and YFV, virus stocks were prepared by infecting sub-confluent Vero or Vero E6 cells at low MOI, collecting supernatants upon the appearance of cytopathic effects following incubation at 37°C in a humidified atmosphere with 5% CO_2_, and clarifying the supernatant through centrifugation at 1,300 rpm for 5 min. DENV-2 and ZIKV were grown for one week on C6/36 cells at 33°C in a humidified atmosphere with 5% CO_2_, and stocks prepared as for the other viruses. For experimental infections, Aag2-AF5 or Aag2-AF256 cells were seeded at 1 × 10^5^ cells per well in 24-well plates and infected one day post-seeding with inoculum made up in culture media containing 2% (v/v) FBS at MOIs as specified in figures. Cells were incubated at room temperature for one hour, washed once with phosphate-buffered saline (PBS), and incubated at 28°C in a humidified atmosphere without CO_2_ following the addition of culture media containing 10% (v/v) FBS.

Viral stock titres were determined by plaque assay titration on Vero cells (CHIKV, JEV, LGTV, MODV, POWV, WNV, YFV and ZIKV) or BHK-21 cells (DENV-2 and RVFV). Briefly, one day post-seeding, confluent cells cultured in 12-well plates were infected with viral inoculum serially diluted in culture media containing 2% FBS (v/v). Following a one-hour incubation, cells were washed in PBS and incubated at 37°C under an overlay comprised of Minimal Eagle’s Medium (MEM) (ThermoFisher Scientific) supplemented with 2% (v/v) FBS and either 0.6% (w/v) colloidal microcrystalline cellulose (Sigma-Aldrich) or 1% (w/v) UltraPure^TM^ Agarose (ThermoFisher Scientific) (ZIKV only). Cells were fixed in formaldehyde and stained with 0.1% (w/v) toluidine blue (ThermoFisher Scientific) to visualise plaques. Where indicated in figures, experimental samples were titrated by 50% tissue culture infectious dose (TCID50) in Vero E6 cells as calculated using the Reed–Muench method. Briefly, confluent cell monolayers were infected with ten-fold serial dilutions of viral inoculum on the day of seeding and were maintained at 37°C for six days post-infection (CHIKV, JEV, POWV, RVFV, WNV and YFV) or nine days post-infection (DENV-2) prior to formaldehyde fixation and staining with 0.1% (w/v) toluidine blue.

### Transient gene silencing

dsRNAs for transient knockdowns were prepared as previously described [15] by amplifying a ∼180-450 nt fragment of the corresponding transcript from Aag2 cells by RT-PCR using primers containing a T7 promoter sequence. dsRNA was generated from the resulting amplicons by *in vitro* transcription using the MEGAshortscript *in vitro* transcription kit (ThermoFisher Scientific) and gel purified before storage in aliquots at -80°C. Primers and dsRNAs against the *REL1A*, *REL2* and *GFP* genes were described previously [15]. Primers for generating dsRNAs against *REL1B* (AAEL006930) were TAATACGACTCACTATAGGGCAGGAAAGGAGGAGCAAACA (prKM80F) and TAATACGACTCACTATAGGGCATAACCCGTCTTGAACGGA (prKM80R). dsRNAs were transfected at the time of seeding into 5 × 10^5^ Aag2-AF5 cells/well (for RT-qPCR and viral infection) or 1.2 × 10^5^ cells/well grown on coverslips (for immunofluorescence microscopy) in 24-well plates at a final concentration of 10 nM using TransIT-insect transient transfection reagent (Cambridge Biosciences, Cambridge, UK) as per manufacturer’s instructions.

### Immune stimulation

NF-κB pathways were stimulated using heat-inactivated *E. coli* DH5α (ThermoFisher Scientific). To prepare bacteria for stimulation, bacteria were cultured overnight at 37°C with shaking in Luria Bertani (LB) medium without antibiotics (Sigma-Aldrich), and titrated on LB agar in 6-well plates. Bacterial cultures were washed in PBS, resuspended in PBS and heat-inactivated at 60°C for 3 h. Inactivation was verified by culture on LB agar prior to storage at -80°C and use in experiments. Aag2-AF5 and Aag2-AF256 cells were stimulated one day post-seeding by replacing the culture medium with fresh medium containing 100 CFU equivalents per cell.

### RNA Purification and RT-qPCR

Total RNA was isolated by column-based extraction using the RNeasy® Mini kit (Qiagen, Hilden, Germany) as per the manufacturer’s instructions following lysis in buffer RLT. cDNA synthesis was performed with 1 µg RNA per 20 µl reaction volume using the SOLIScript® RT cDNA synthesis MIX kit (Solis BioDyne, Tartu, Estonia) as per the manufacturer’s instructions with the addition of 50 ng random hexamers (ThermoFisher Scientific). The thermocycler conditions were 10 min primer annealing at 25°C, followed by 50 min at 50°C and reaction termination at 85°C for 5 min. Intracellular viral RNA was quantified relative to the 40S ribosomal protein S7 (*RPS7*) mRNA (2^-ΔCt^) by two-step RT-qPCR performed using Fast SYBR Green (Applied Biosystems, Waltham, MA USA) as per the manufacturer’s instructions using 1 μl cDNA and 100 nM primers per 10 µl reaction, as detailed in Table 1. Thermocycling was performed at 95°C for 20 s, followed by 40 cycles of 95°C for 1 s and 60°C for 20 s, as per the manufacturer’s instructions.

**Table 1.**
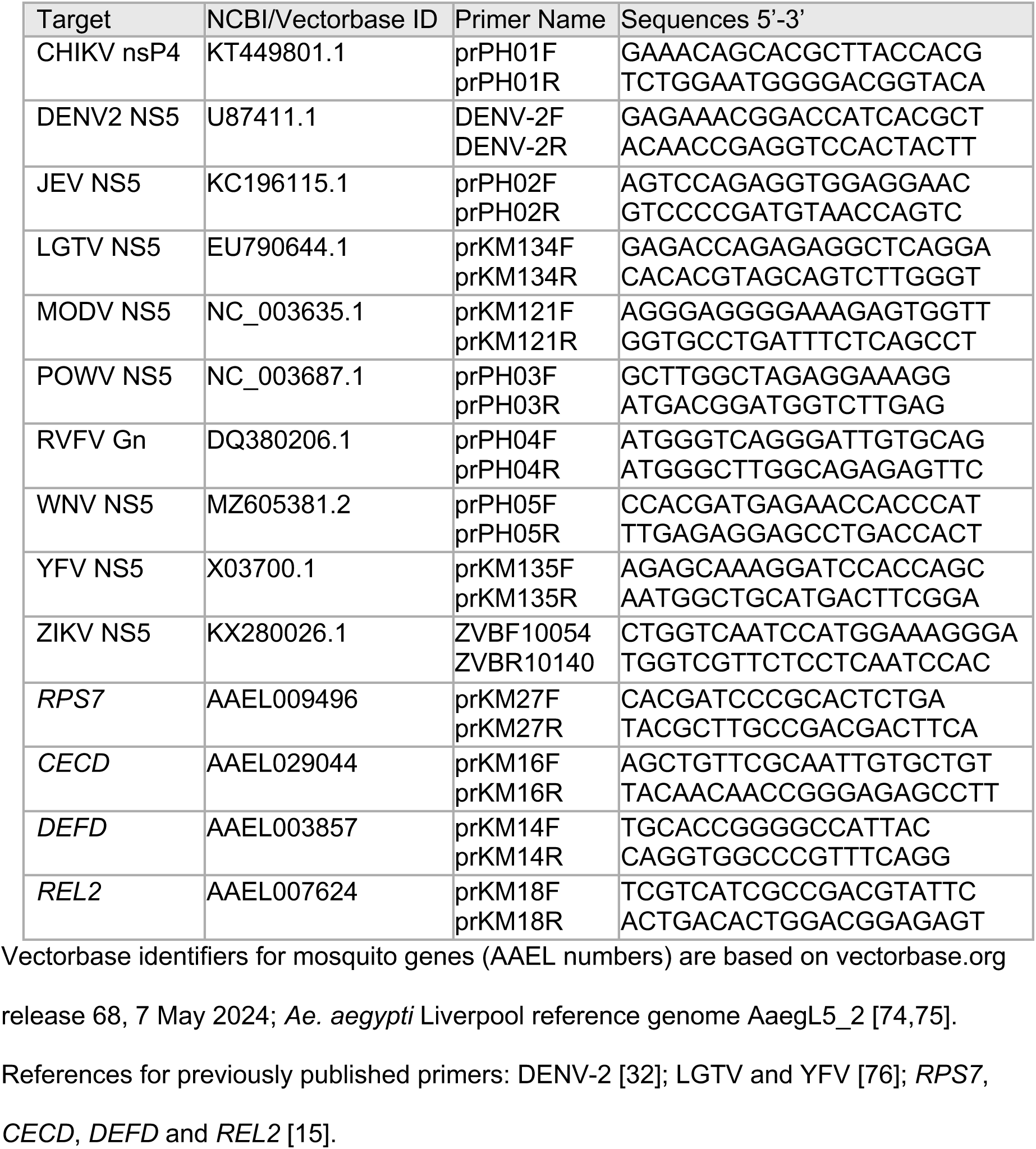
qPCR primers used in this study.

### Confocal immunofluorescence microscopy

Ankyrons for the detection of *Ae. aegypti* NF-κB proteins were produced by ProImmune (Oxford, UK). Briefly, recombinant protein fragments corresponding to Rel1A (Uniprot ID Q171A6) residues ∼50-220, Rel1B (Q174E0) residues ∼160-265 or Rel2 (Q171I8) residues ∼130-320 were used to identify NF-κB-binding Ankyrons from ProImmune’s Teralibrary of one trillion variants. Resulting Ankyrons were tested in our hands for detection of NF-κB proteins by immunofluorescence microscopy, and specificity was confirmed using Aag2-AF256 Rel2 knockout cells or dsRNA-mediated transient knockdowns of Rel1A or Rel1B. ProImmune Ankyron clone IDs for the data shown here are BT40163 (Rel1A), BT40219 (Rel1B) and BT40178 (Rel2).

For immunofluorescence microscopy, cells were fixed three days post-seeding with 4% paraformaldehyde for 20 min and washed three times in PBS for 5 min. Cells were then blocked and permeabilised in a solution of 5% bovine serum albumin (BSA) and 0.3% Triton X-100 (both Sigma-Aldrich) in PBS for one hour. Cells were incubated with V5-tagged Ankyrons at a final concentration of 0.05 mg/ml, then mouse anti-SV5 antibody (1:50 dilution, ab27671, Abcam, Cambridge, UK) and goat anti-mouse AlexaFluor^TM^ 488 fluorescent secondary antibody (1:5000, A-11029, ThermoFisher Scientific) at room temperature for one hour each in a solution of 1% BSA, 0.3% Triton X-100 in PBS, with three five-min PBS washes between Ankyron/antibody incubations. Slides were mounted in SlowFade^TM^ Gold Antifade Mountant with 4′,6-diamidino-2-phenylindole (DAPI) (ThermoFisher Scientific). Images were captured at 63× magnification using a Leica STELLARIS 5 confocal inverted microscope and LAS X Life Science Microscope Software (Leica Microsystems, Wetzlar, Germany).

### Data visualisation and statistical analysis

Numerical data were analysed using Microsoft Excel (Microsoft Corporation, Redmund, WA USA) and GraphPad Prism version 10.5.0 (Dotmatics, Boston, MA USA). All statistical analyses were performed in GraphPad Prism, using two-way ANOVA with Tukey’s correction, apart from Fig 6Aiii and 6Ciii, which were analysed using Student’s *t*-test with Welch’s correction, and Fig 6Biii, which was analysed by one-way ANOVA with Dunnett’s correction. Figures were prepared using Adobe Illustrator and Adobe Photoshop (Adobe Systems, San Jose, CA USA). Microscopy images were cropped, annotated and modified to optimise brightness and contrast only, with all concurrently acquired images being edited at the same time to avoid the appearance of artefacts.

## Acknowledgements

We acknowledge the support of the Bioimaging, Cell Culture and CL3 Virology facilities at The Pirbright Institute, whose resources were instrumental in facilitating this research.

## Author Credit

Conceptualization: ACF, KM

Data curation: PH, KM

Formal analysis: PH, YPC, TAR, KM

Funding acquisition: KM

Investigation: PH, YPC, RA, TAR, ACF, VK, LEW, KM

Methodology: PH, YPC, RA, TAR, ACF, VK, KM

Resources: ID, TM Supervision: ADD, AFS, KM

Visualization: PH, RA, TAR, VK, KM

Writing – original draft: PH, KM

Writing – review & editing: all authors

## References

1. Jones KE, Patel NG, Levy MA, Storeygard A, Balk D, Gittleman JL, et al. Global trends in emerging infectious diseases. Nature. 2008;451: 990–993. doi:10.1038/nature06536

2. Ellis BR, Wilcox BA. The ecological dimensions of vector-borne disease research and control. Cadernos de saude publica. 2009;25 Suppl 1: S155–67.

3. Kramer LD. Complexity of virus-vector interactions. Current Opinion in Virology. 2016;21: 81–86. doi:10.1016/j.coviro.2016.08.008

4. Ahmad NA, Mancini M-V, Ant TH, Martinez J, Kamarul GMR, Nazni WA, et al. Wolbachia strain w AlbB maintains high density and dengue inhibition following introduction into a field population of Aedes aegypti. Philosophical Transactions Royal Soc B Biological Sci. 2020;376: 20190809. doi:10.1098/rstb.2019.0809

5. Sinkins SP. Wolbachia and arbovirus inhibition in mosquitoes. Future Microbiol. 2013;8: 1249–1256. doi:10.2217/fmb.13.95

6. O’Neill SL. The Use of Wolbachia by the World Mosquito Program to Interrupt Transmission of Aedes aegypti Transmitted Viruses. Adv Exp Med Biol. 2018;1062: 355–360. doi:10.1007/978-981-10-8727-1_24

7. Reitmayer CM, Levitt E, Basu S, Atkinson B, Fragkoudis R, Merits A, et al. Mimicking superinfection exclusion disrupts alphavirus infection and transmission in the yellow fever mosquito Aedes aegypti. Proc Natl Acad Sci United States Am. 2023;120: e2303080120. doi:10.1073/pnas.2303080120

8. Buchman A, Gamez S, Li M, Antoshechkin I, Li H-H, Wang H-W, et al. Broad dengue neutralization in mosquitoes expressing an engineered antibody. Plos Pathog. 2020;16: e1008103. doi:10.1371/journal.ppat.1008103

9. Buchman A, Gamez S, Li M, Antoshechkin I, Li H-H, Wang H-W, et al. Engineered resistance to Zika virus in transgenic Aedes aegypti expressing a polycistronic cluster of synthetic small RNAs. P Natl Acad Sci Usa. 2019;116: 3656–3661. doi:10.1073/pnas.1810771116

10. Viglietta M, Bellone R, Blisnick AA, Failloux A-B. Vector Specificity of Arbovirus Transmission. Front Microbiol. 2021;12: 773211. doi:10.3389/fmicb.2021.773211

11. Merkling SH, Rij RP van. Beyond RNAi: Antiviral defense strategies in Drosophila and mosquito. J Insect Physiol. 2013;59: 159–170. doi:10.1016/j.jinsphys.2012.07.004

12. Bronkhorst AW, Rij RP van. The long and short of antiviral defense: small RNA-based immunity in insects. Current Opinion in Virology. 2014;7: 19–28. doi:10.1016/j.coviro.2014.03.010

13. Liu W-L, Hsu C-W, Chan S-P, Yen P-S, Su MP, Li J-C, et al. Transgenic refractory Aedes aegypti lines are resistant to multiple serotypes of dengue virus. Sci Rep. 2021;11: 23865. doi:10.1038/s41598-021-03229-4

14. Cheung YP, Park S, Pagtalunan J, Maringer K. The antiviral role of NF-κB-mediated immune responses and their antagonism by viruses in insects. J Gen Virol. 2022;103. doi:10.1099/jgv.0.001741

15. Russell TA, Ayaz A, Davidson AD, Fernandez-Sesma A, Maringer K. Imd pathway-specific immune assays reveal NF-κB stimulation by viral RNA PAMPs in Aedes aegypti Aag2 cells. Plos Neglect Trop D. 2021;15: e0008524. doi:10.1371/journal.pntd.0008524

16. Angleró-Rodríguez YI, Tikhe CV, Kang S, Dimopoulos G. Aedes aegypti Toll pathway is induced through dsRNA sensing in endosomes. Dev Comp Immunol. 2021;122: 104138. doi:10.1016/j.dci.2021.104138

17. Prince BC, Chan K, Rückert C. Elucidating the role of dsRNA sensing and Toll6 in antiviral responses of Culex quinquefasciatus cells. Front Cell Infect Microbiol. 2023;13: 1251204. doi:10.3389/fcimb.2023.1251204

18. Fragkoudis R, Chi Y, Siu RWC, Barry G, Attarzadeh-Yazdi G, Merits A, et al. Semliki Forest virus strongly reduces mosquito host defence signaling. Insect Molecular Biology. 2008;17: 647–656. doi:10.1111/j.1365-2583.2008.00834.x

19. Xiao X, Liu Y, Zhang X, Wang J, Li Z, Pang X, et al. Complement-Related Proteins Control the Flavivirus Infection of Aedes aegypti by Inducing Antimicrobial Peptides. PLOS Pathog. 2014;10: e1004027. doi:10.1371/journal.ppat.1004027

20. Chowdhury A, Modahl CM, Tan ST, Xiang BWW, Missé D, Vial T, et al. JNK pathway restricts DENV2, ZIKV and CHIKV infection by activating complement and apoptosis in mosquito salivary glands. Plos Pathog. 2020;16: e1008754. doi:10.1371/journal.ppat.1008754

21. Shi Z-K, Wen D, Chang M-M, Sun X-M, Wang Y-H, Cheng C-H, et al. Juvenile Hormone-Sensitive Ribosomal Activity Enhances Viral Replication in Aedes aegypti. Msystems. 2021; e0119020. doi:10.1128/msystems.01190-20

22. Weng S-C, Li H-H, Li J-C, Liu W-L, Chen C-H, Shiao S-H. A Thioester-Containing Protein Controls Dengue Virus Infection in Aedes aegypti Through Modulating Immune Response. Front Immunol. 2021;12: 670122. doi:10.3389/fimmu.2021.670122

23. Waterhouse RM, Kriventseva EV, Meister S, Xi Z, Alvarez KS, Bartholomay LC, et al. Evolutionary dynamics of immune-related genes and pathways in disease-vector mosquitoes. Science. 2007;316: 1738–1743. doi:10.1126/science.1139862

24. Tzou P, Ohresser S, Ferrandon D, Capovilla M, Reichhart JM, Lemaitre B, et al. Tissue-specific inducible expression of antimicrobial peptide genes in Drosophila surface epithelia. Immunity. 2000;13: 737–748.

25. Silverman N, Zhou R, Erlich RL, Hunter M, Bernstein E, Schneider D, et al. Immune activation of NF-kappaB and JNK requires Drosophila TAK1. The Journal of Biological Chemistry. 2003;278: 48928–48934. doi:10.1074/jbc.m304802200

26. Delaney JR, Stöven S, Uvell H, Anderson KV, Engstrom Y, Mlodzik M. Cooperative control of Drosophila immune responses by the JNK and NF-kappaB signaling pathways. EMBO J. 2006;25: 3068–3077. doi:10.1038/sj.emboj.7601182

27. Chowdhury M, Zhang J, Xu X-X, He Z, Lu Y, Liu X-S, et al. An in vitro study of NF-κB factors cooperatively in regulation of Drosophila melanogaster antimicrobial peptide genes. Dev Comp Immunol. 2019;95: 50–58. doi:10.1016/j.dci.2019.01.017

28. Gregorio ED, Spellman PT, Tzou P, Rubin GM, Lemaitre B. The Toll and Imd pathways are the major regulators of the immune response in Drosophila. The EMBO Journal. 2002;21: 2568–2579. doi:10.1093/emboj/21.11.2568

29. Bruurs LJM, Müller M, Schipper JG, Rabouw HH, Boersma S, Kuppeveld FJM van, et al. Antiviral responses are shaped by heterogeneity in viral replication dynamics. Nat Microbiol. 2023;8: 2115–2129. doi:10.1038/s41564-023-01501-z

30. Varjak M, Maringer K, Watson M, Sreenu VB, Fredericks AC, Pondeville E, et al. Aedes aegypti Piwi4 Is a Noncanonical PIWI Protein Involved in Antiviral Responses. Duprex WP, editor. mSphere. 2017;2: e00144–17. doi:10.1128/msphere.00144-17

31. Scherer C, Knowles J, Sreenu VB, Fredericks AC, Fuss J, Maringer K, et al. An Aedes aegypti-Derived Ago2 Knockout Cell Line to Investigate Arbovirus Infections. Viruses. 2021; 1066.

32. Fredericks AC, Russell TA, Wallace LE, Davidson AD, Fernandez-Sesma A, Maringer K. Aedes aegypti (Aag2)-derived clonal mosquito cell lines reveal the effects of pre-existing persistent infection with the insect-specific bunyavirus Phasi Charoen-like virus on arbovirus replication. PLOS Neglect Trop D. 2019;13: e0007346. doi:10.1371/journal.pntd.0007346

33. Shin SW, Kokoza V, Ahmed A, Raikhel AS. Characterization of three alternatively spliced isoforms of the Rel/NF-κB transcription factorRelish from the mosquito Aedes aegypti. Proceedings of the National Academy of Sciences. 2002;99: 9978–9983. Available: http://www.pnas.org/content/99/15/9978.full.pdf#page=1&view=FitH

34. Luplertlop N, Surasombatpattana P, Patramool S, Dumas E, Wasinpiyamongkol L, Saune L, et al. Induction of a Peptide with Activity against a Broad Spectrum of Pathogens in the Aedes aegypti Salivary Gland, following Infection with Dengue Virus. PLOS Pathog. 2011;7: e1001252. doi:10.1371/journal.ppat.1001252.s003

35. Almire F, Terhzaz S, Terry S, McFarlane M, Gestuveo RJ, Szemiel AM, et al. Sugar feeding protects against arboviral infection by enhancing gut immunity in the mosquito vector Aedes aegypti. PLoS Pathog. 2021;17: e1009870. doi:10.1371/journal.ppat.1009870

36. Barletta ABF, Nascimento-Silva MCL, Talyuli OAC, Oliveira JHM, Pereira LOR, Oliveira PL, et al. Microbiota activates IMD pathway and limits Sindbis infection in Aedes aegypti. Parasites & Vectors. 2017;10: 103. doi:10.1186/s13071-017-2040-9

37. Zakovíc S, Rivera GE, Gomaid R, Martinez CG, Kappler C, Marois E, et al. The major role of the REL2/NF-κB pathway in the regulation of midgut bacterial homeostasis in the malaria vector Anopheles gambiae. bioRxiv. 2025; 2025.03.14.643338. doi:10.1101/2025.03.14.643338

38. Ramirez JL, Souza-Neto J, Cosme RT, Rovira J, Ortiz A, Pascale JM, et al. Reciprocal Tripartite Interactions between the Aedes aegypti Midgut Microbiota, Innate Immune System and Dengue Virus Influences Vector Competence. O’Neill SL, editor. PLoS Neglected Tropical Diseases. 2012;6: e1561. doi:10.1371/journal.pntd.0001561.s004

39. Song X, Zhou H, Wang J. Cell wall components of gut commensal bacteria stimulate peritrophic matrix formation in malaria vector mosquitoes through activation of the IMD pathway. PLOS Biol. 2025;23: e3002967. doi:10.1371/journal.pbio.3002967

40. Accoti A, Becker M, Abu AEI, Vulcan J, Jun R, Widen SG, et al. Dehydration-induced Ae-Aper50 regulates midgut infection in Aedes aegypti mosquitoes. mBio. 2025;16: e01207–24. doi:10.1128/mbio.01207-24

41. Talyuli OAC, Oliveira JHM, Bottino-Rojas V, Silveira GO, Alvarenga PH, Barletta ABF, et al. The Aedes aegypti peritrophic matrix controls arbovirus vector competence through HPx1, a heme–induced peroxidase. PLOS Pathog. 2023;19: e1011149. doi:10.1371/journal.ppat.1011149

42. Cai H, Holleufer A, Simonsen B, Schneider J, Lemoine A, Gad HH, et al. 2’3’-cGAMP triggers a STING- and NF-κB-dependent broad antiviral response in Drosophila. Sci Signal. 2020;13. doi:10.1126/scisignal.abc4537

43. Cai H, Holleufer A, Simonsen B, Schneider J, Lemoine A, Gad H-H, et al. 2’3’-cGAMP triggers a STING and NF-κB dependent broad antiviral response in Drosophila. bioRxiv. 2019; 1–33. doi:10.1101/852319

44. Slavik KM, Morehouse BR, Ragucci AE, Zhou W, Ai X, Chen Y, et al. cGAS-like receptors sense RNA and control 3′2′-cGAMP signaling in Drosophila. Nature. 2021; 1–8. doi:10.1038/s41586-021-03743-5

45. Holleufer A, Winther KG, Gad HH, Ai X, Chen Y, Li L, et al. Two cGAS-like receptors induce antiviral immunity in Drosophila. Nature. 2021; 1–7. doi:10.1038/s41586-021-03800-z

46. Wu X, Wu F-H, Wang X, Wang L, Siedow JN, Zhang W, et al. Molecular evolutionary and structural analysis of the cytosolic DNA sensor cGAS and STING. Nucleic Acids Res. 2014;42: 8243–8257. doi:10.1093/nar/gku569

47. Paradkar PN, Duchemin J-B, Voysey R, Walker PJ. Dicer-2-Dependent Activation of Culex Vago Occurs via the TRAF-Rel2 Signaling Pathway. Olson KE, editor. PLOS Neglect Trop D. 2014;8: e2823. doi:10.1371/journal.pntd.0002823

48. Sim S, Dimopoulos G. Dengue Virus Inhibits Immune Responses in Aedes aegypti Cells. PLoS ONE. 2010;5: e10678. doi:10.1371/journal.pone.0010678.s006

49. Pompon J, Manuel M, Ng GK, Wong B, Shan C, Manokaran G, et al. Dengue subgenomic flaviviral RNA disrupts immunity in mosquito salivary glands to increase virus transmission. PLoS pathogens. 2017;13: e1006535. doi:10.1371/journal.ppat.1006535

50. Méndez Y, Pacheco C, Herrera F. Inhibition of defensin A and cecropin A responses to dengue virus 1 infection in Aedes aegypti. Biomédica. 2020;41: 161–167. doi:10.7705/biomedica.5491

51. Paradkar PN, Trinidad L, Voysey R, Duchemin J-B, Walker PJ. Secreted Vago restricts West Nile virus infection in Culex mosquito cells by activating the Jak-STAT pathway. PNAS. 2012; 1–6. doi:10.1073/pnas.1205231109/-/dcsupplemental/sapp.docx

52. Deddouche S, Matt N, Budd A, Mueller S, Kemp C, Galiana-Arnoux D, et al. The DExD/H-box helicase Dicer-2 mediates the induction of antiviral activity in drosophila. Nat Immunol. 2008;9: 1425–1432. doi:10.1038/ni.1664

53. Couderc E, Crist AB, Daron J, Varet H, Hout FAH van, Miesen P, et al. A Vago-like gene enhances dengue and Zika virus dissemination in Aedes aegypti. bioRxiv. 2024; 2024.07.01.601473. doi:10.1101/2024.07.01.601473

54. Zaritsky LA, Bedsaul JR, Zoon KC. Virus Multiplicity of Infection Affects Type I Interferon Subtype Induction Profiles and Interferon-Stimulated Genes. J Virol. 2015;89: 11534–11548. doi:10.1128/jvi.01727-15

55. Mehta D, Chaudhary S, Sunil S. Oxidative stress governs mosquito innate immune signalling to reduce chikungunya virus infection in Aedes-derived cells. J Gen Virol. 2024;105. doi:10.1099/jgv.0.001966

56. Carissimo G, Pondeville E, McFarlane M, Dietrich I, Mitri C, Bischoff E, et al. Antiviral immunity of Anopheles gambiae is highly compartmentalized, with distinct roles for RNA interference and gut microbiota. Proceedings of the National Academy of Sciences. 2014;112: E176–E185. doi:10.1073/pnas.1412984112

57. Waldock J, Olson KE, Christophides GK. Anopheles gambiae Antiviral Immune Response to Systemic O’nyong-nyong Infection. Traub-Csekö YM, editor. PLoS Neglected Tropical Diseases. 2012;6: e1565. doi:10.1371/journal.pntd.0001565.s004

58. Laureti M, Lee R-X, Bennett A, Wilson LA, Sy VE, Kohl A, et al. Rift Valley Fever Virus Primes Immune Responses in Aedes aegypti Cells. Pathogens. 2023;12: 563. doi:10.3390/pathogens12040563

59. Núñez AI, Esteve-Codina A, Gómez-Garrido J, Brustolin M, Talavera S, Berdugo M, et al. Alteration in the Culex pipiens transcriptome reveals diverse mechanisms of the mosquito immune system implicated upon Rift Valley fever phlebovirus exposure. PLoS Neglected Trop Dis. 2020;14: e0008870. doi:10.1371/journal.pntd.0008870

60. Liu K, Xiao C, Xi S, Hameed M, Wahaab A, Shao D, et al. Mosquito defensins enhance Japanese encephalitis virus infection by facilitating virus adsorption and entry within mosquito. 2020; 1–58. doi:10.1101/2020.04.28.065904

61. Charlier N, Davidson A, Dallmeier K, Molenkamp R, Clercq ED, Neyts J. Replication of not-known-vector flaviviruses in mosquito cells is restricted by intracellular host factors rather than by the viral envelope proteins. Journal of General Virology. 2010;91: 1693–1697. doi:10.1099/vir.0.019851-0

62. Junglen S, Korries M, Grasse W, Wieseler J, Kopp A, Hermanns K, et al. Host Range Restriction of Insect-Specific Flaviviruses Occurs at Several Levels of the Viral Life Cycle. Randall G, editor. 2017;2: e00375–16. doi:10.1128/msphere.00375-16

63. Tree MO, McKellar DR, Kieft KJ, Watson AM, Ryman KD, Conway MJ. Insect-specific flavivirus infection is restricted by innate immunity in the vertebrate host. Virology. 2016;497: 81–91. doi:10.1016/j.virol.2016.07.005

64. Ramirez JL, Dimopoulos G. The Toll immune signaling pathway control conserved anti-dengue defenses across diverse Ae. aegypti strains and against multiple dengue virus serotypes. Dev Comp Immunol. 2010;34: 625–629. doi:10.1016/j.dci.2010.01.006

65. Xi Z, Ramirez JL, Dimopoulos G. The Aedes aegypti toll pathway controls dengue virus infection. PLOS Pathog. 2008;4: e1000098. doi:10.1371/journal.ppat.1000098

66. Brackney DE, Scott JC, Sagawa F, Woodward JE, Miller NA, Schilkey FD, et al. C6/36 Aedes albopictus Cells Have a Dysfunctional Antiviral RNA Interference Response. O’Neill SL, editor. PLoS Neglected Tropical Diseases. 2010;4: e856. doi:10.1371/journal.pntd.0000856.t001

67. Lawrie CH, Uzcátegui NY, Armesto M, Bell-Sakyi L, Gould EA. Susceptibility of mosquito and tick cell lines to infection with various flaviviruses. Méd Vet Èntomol. 2004;18: 268–274. doi:10.1111/j.0269-283x.2004.00505.x

68. Varjak M, Donald CL, Mottram TJ, Sreenu VB, Merits A, Maringer K, et al. Characterization of the Zika virus induced small RNA response in Aedes aegypti cells. Olson KE, editor. PLOS Neglect Trop D. 2017;11: e0006010. doi:10.1371/journal.pntd.0006010

69. Hoermann A, Habtewold T, Selvaraj P, Corsano GD, Capriotti P, Inghilterra MG, et al. Gene drive mosquitoes can aid malaria elimination by retarding Plasmodium sporogonic development. Sci Adv. 2022;8: eabo1733. doi:10.1126/sciadv.abo1733

70. Merkling SH, Crist AB, Henrion-Lacritick A, Frangeul L, Couderc E, Gausson V, et al. Multifaceted contributions of Dicer2 to arbovirus transmission by Aedes aegypti. Cell Rep. 2023;42: 112977. doi:10.1016/j.celrep.2023.112977

71. Maringer K. Re-evaluating the mosquito RNAi pathway’s influence on arbovirus transmission. Trends Parasitol. 2023. doi:10.1016/j.pt.2023.09.005

72. Samuel GH, Pohlenz T, Dong Y, Coskun N, Adelman ZN, Dimopoulos G, et al. RNA interference is essential to modulating the pathogenesis of mosquito-borne viruses in the yellow fever mosquito Aedes aegypti. Proc National Acad Sci. 2023;120: e2213701120. doi:10.1073/pnas.2213701120

73. Bassett AR, Tibbit C, Ponting CP, Liu J-L. Mutagenesis and homologous recombination in Drosophila cell lines using CRISPR/Cas9. Biology open. 2014;3: 42–49. doi:10.1242/bio.20137120

74. Giraldo-Calderón GI, Emrich SJ, Maccallum RM, Maslen G, Dialynas E, Topalis P, et al. VectorBase: an updated bioinformatics resource for invertebrate vectors and other organisms related with human diseases. Nucleic Acids Research. 2015;43: D707–13. doi:10.1093/nar/gku1117

75. Matthews BJ, Dudchenko O, Kingan SB, Koren S, Antoshechkin I, Crawford JE, et al. Improved reference genome of Aedes aegypti informs arbovirus vector control. Nature. 2018;563: 501–507. doi:10.1038/s41586-018-0692-z

76. Taguwa S, Maringer K, Li X, Bernal-Rubio D, Rauch JN, Gestwicki JE, et al. Defining Hsp70 Subnetworks in Dengue Virus Replication Reveals Key Vulnerability in Flavivirus Infection. Cell. 2015;163: 1108–1123. doi:10.1016/j.cell.2015.10.046

